# *HaVoC*, a bioinformatic pipeline for reference-based consensus assembly and lineage assignment for SARS-CoV-2 sequences

**DOI:** 10.1101/2021.02.12.431018

**Authors:** Phuoc Truong Nguyen, Ilya Plyusnin, Tarja Sironen, Olli Vapalahti, Ravi Kant, Teemu Smura

## Abstract

**Background:** SARS-CoV-2 related research has increased in importance worldwide since December 2019. Several new variants of SARS-CoV-2 have emerged globally, of which the most notable and concerning currently are the UK variant B.1.1.7, the South African variant B1.351 and the Brazilian variant P.1. Detecting and monitoring novel variants is essential in SARS-CoV-2 surveillance. While there are several tools for assembling virus genomes and performing lineage analyses to investigate SARS-CoV-2, each is limited to performing singular or a few functions separately.

**Results:** Due to the lack of publicly available pipelines, which could perform fast reference-based assemblies on raw SARS-CoV-2 sequences in addition to identifying lineages to detect variants of concern, we have developed an open source bioinformatic pipeline called *HaVoC* (Helsinki university Analyzer for Variants Of Concern). *HaVoC* can reference assemble raw sequence reads and assign the corresponding lineages to SARS-CoV-2 sequences.

**Conclusions:** *HaVoC* is a pipeline utilizing several bioinformatic tools to perform multiple necessary analyses for investigating genetic variance among SARS-CoV-2 samples. The pipeline is particularly useful for those who need a more accessible and fast tool to detect and monitor the spread of SARS-CoV-2 variants of concern during local outbreaks. *HaVoC* is currently being used in Finland for monitoring the spread of SARS-CoV-2 variants. *HaVoC* user manual and source code are available at https://www.helsinki.fi/en/projects/havoc and https://bitbucket.org/auto_cov_pipeline/havoc, respectively.

## Background

Emerging pathogens pose a continuous threat to mankind, as exemplified by the Ebola virus epidemic in West Africa in 2014 [1], Zika virus pandemic in 2015 [2], and the ongoing Coronavirus disease 2019 (COVID-19) pandemic. These viruses are zoonotic, i.e. have crossed species barriers from animals to humans, alike the majority of emerging human pathogens [3, 4]. The likelihood of this host switching is enhanced by several factors, e.g. global movement of people and animals, environmental changes, increased proximity of humans, wildlife and livestock, and population expansion into new environments [5].

The mutation and evolution rate of RNA viruses is considerably higher than their hosts, which is advantageous for viral adaptation. Mutations in the viral genome are most of the time silent or, if affecting phenotype, related to attenuation, although mutations can also lead to more pathogenic strains. A new virus variant may have one or more mutations that separate it from the wild-type virus already circulating among the general population.

Coronaviruses (family *Coronaviridae*) are enveloped single-stranded RNA viruses, which cause respiratory, enteric, hepatic, and neurological diseases of a broad spectrum of severity among different animals and humans. Severe acute respiratory syndrome coronavirus 2 (SARS-CoV-2), a novel evolutionary divergent virus responsible for the present pandemic, has devastated societies and economies globally. The SARS-CoV-2 pandemic has already infected more than 100 million people in 221 countries, causing over 2.2 million global deaths as of 3^rd^ February 2021 [6]. In autumn 2020, a new variant of SARS-CoV-2 known as 20B/501Y.V1 (B.1.1.7) was detected in south-eastern England, Wales, and Scotland [7]. This variant has since spread globally to more than 80 countries. The variant has undergone 23 mutations with 13-nonsynonymous mutations, four amino acid deletions, and six synonymous mutations making the virus more transmissible [8]. Another variant 20C/501Y.V2 (B.1.351) was detected in South Africa which was genetically distant from the UK 20B/501Y.V1 variant [9]. This South African variant with its two mutations in the receptor-binding motif that mainly forms the interface with the human ACE2 receptor has also been widely spreading to circulate globally. It has been noticed that some existing vaccines against SARS-CoV-2 are less effective against the 20C/501Y.V2 variant [10–12]. A third variant being closely monitored is P.1 detected first in Brazil [13]. Interestingly, all these three variants have a mutation in the receptor binding domain (RBD) of the spike protein at position 501, where the amino acid asparagine (N) has been replaced with tyrosine (Y) enabling specific PCR to detect the N501Y mutation [14].

As more transmissible coronavirus variants are circulating worldwide, the role of researchers and technology specialists in controlling the pandemic has received more emphasis. The surveillance of virus variants by sequencing the SARS-CoV-2 genomes would provide a fast way to monitor variants and their spread, however, there are only few publicly available methods for quick reference-based consensus assembly and lineage assignment for SARS-CoV-2 samples. For this purpose, we have developed a simple pipeline, called *HaVoC* (Helsinki university Analyzer for Variants Of Concern), for quick reference-based consensus assembly and lineage assignment for SARS-CoV-2 samples. This will provide the end user a quick and accessible method of variant identification and monitoring. The pipeline was developed to be run on Unix/Linux operating systems, and thus can also be used in remote servers, e.g. CSC – IT Center for Science, Finland.

## Implementation

*HaVoC* consists of a single shell script, which performs reference-based consensus assemblies to query SARS-CoV-2 fastq sequence libraries and assigns lineages to them individually in succession. The script can be started by typing the following line into your command line terminal:

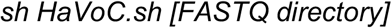

The computing of consensus sequences starts with the tool detecting FASTQ files generated via paired end sequencing in a given input directory and checking that each query FASTQ file has its corresponding counterpart, i.e. mates file. The names of the files are modified to be more concise, e.g. Query-Seq:1_X123_Y000_R1_000.fastq.gz to Query-Seq:1_R1.fastq.gz. The pipeline accepts FASTQ files both in gzipped and uncompressed format.

For the analyses, the user can choose which bioinformatic tools to utilize. This can be done by typing the tool wanted (*tools_prepro, tools_aligner* and *tools_sam*) within the options section in the beginning of the script file. For example, if the user wants to deploy Trimmomatic to pre-process FASTQ files, the following line can be changed as follows:

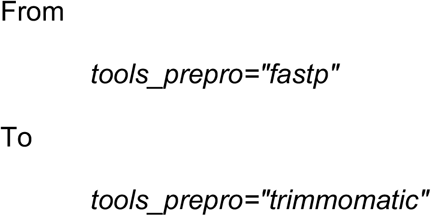

Other options include the number of threads, minimum coverage below which a region is masked (*min_coverage*), and whether to run Pangolin to assign lineages to the consensus genome (*run_pangolin*). An additional option allows *HaVoC* to be run in the CSC servers (*run_in_csc*).

The pre-alignment quality control, e.g. removing and trimming low quality reads and bases, removing adapter sequences, can be done with either fastp [15] or Trimmomatic [16]. The reads are then aligned to a reference genome of SARS-CoV-2 isolate Wuhan-Hu-1 (Genbank accession code: NC_045512.2) with BWA-MEM [17] or Bowtie 2 [18]. The resulting SAM and BAM files are processed (includes sorting, filling in mate coordinates, marking duplicate alignments, and indexing reads) with Sambamba [19] or Samtools [20] and the low coverage regions are masked with BEDtools [21]. After masking a variant call is done with Lofreq [22] before computing the consensus sequence via BCFtools of Samtools [20]. Finally, the consensus sequence is analyzed with pangolin [23] to assign a lineage. The whole process is depicted in figure 1.

**Fig. 1.**
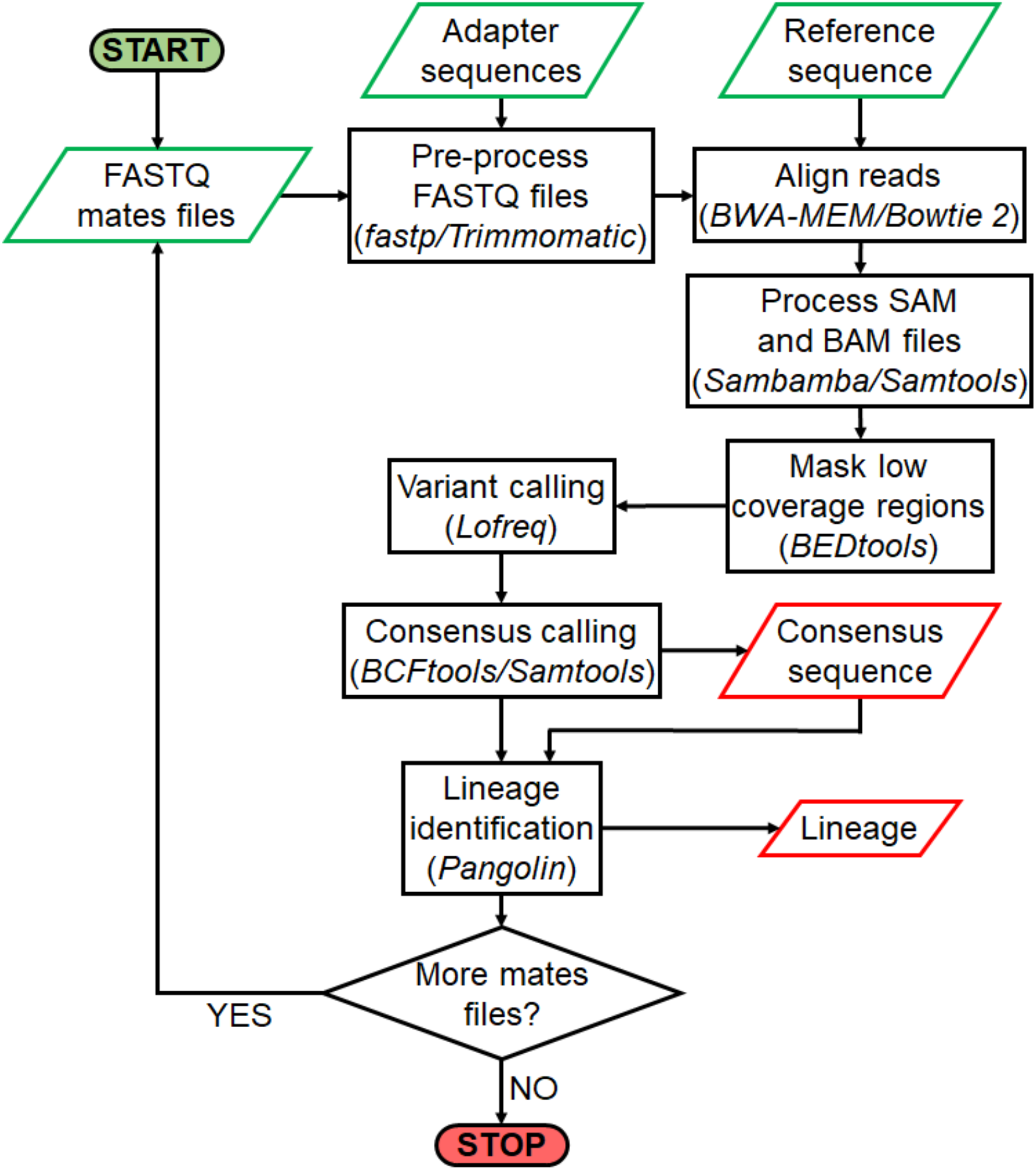
Flowchart describing processes and steps performed by *HaVoC* pipeline. The pipeline constructs consensus sequences from all FASTQ files in an input directory and then compares the resulting sequences to other established SARS-CoV-2 genomes to assign them the most likely lineages. The pipeline requires a FASTA file of adapter sequences for FASTQ pre-processing and a reference genome of SARS-CoV-2 in a separate FASTA file. The adapter file is not required when running the pipeline with fastp option. Input files are highlighted in green and the outputs in red.

### Usage example

We are going to demonstrate a common use case for *HaVoC* with FASTQ files containing reads for SARS-CoV-2 sequences, provided by the Viral zoonoses research unit at University of Helsinki, Finland. The test files within the Example_FASTQs folder contain paired-end FASTQ files for the UK variant (UK-variant-1) and the South African variant (S-Africa-variant-1). To analyse these example files, the aforementioned command needs to be deployed as follows:

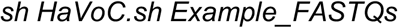

## Results

The FASTQ files are processed and analyzed with the default options utilizing faster bioinformatic tools (fastp, BWA-MEM and Sambamba) in ca. 2–4 minutes, depending on the performance of the platform (local or server). After *HaVoc* has finished the analyses, each FASTQ file is moved to their respective result folders within the FASTQ directory. Each result folder contains a FASTA file for the consensus sequence (e.g. UK-variant-1_consensus.fa) and a CSV file with the lineage information produced by pangolin (e.g. UK-variant-1_pangolin_lineage.csv). In addition to these main result files, each directory contains the original FASTQ files, BAM files (original, indexed and sorted), variant call files (VCF) with mutation data, BED file used for masking regions, and fastp report files with the results of FASTQ processing. The resulting directory and file structure with the example files will look as follows:

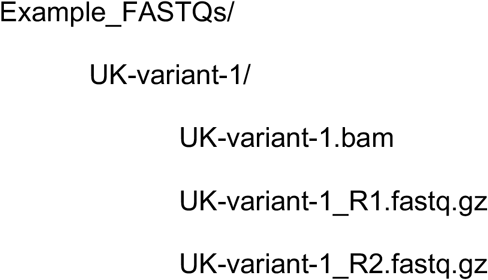

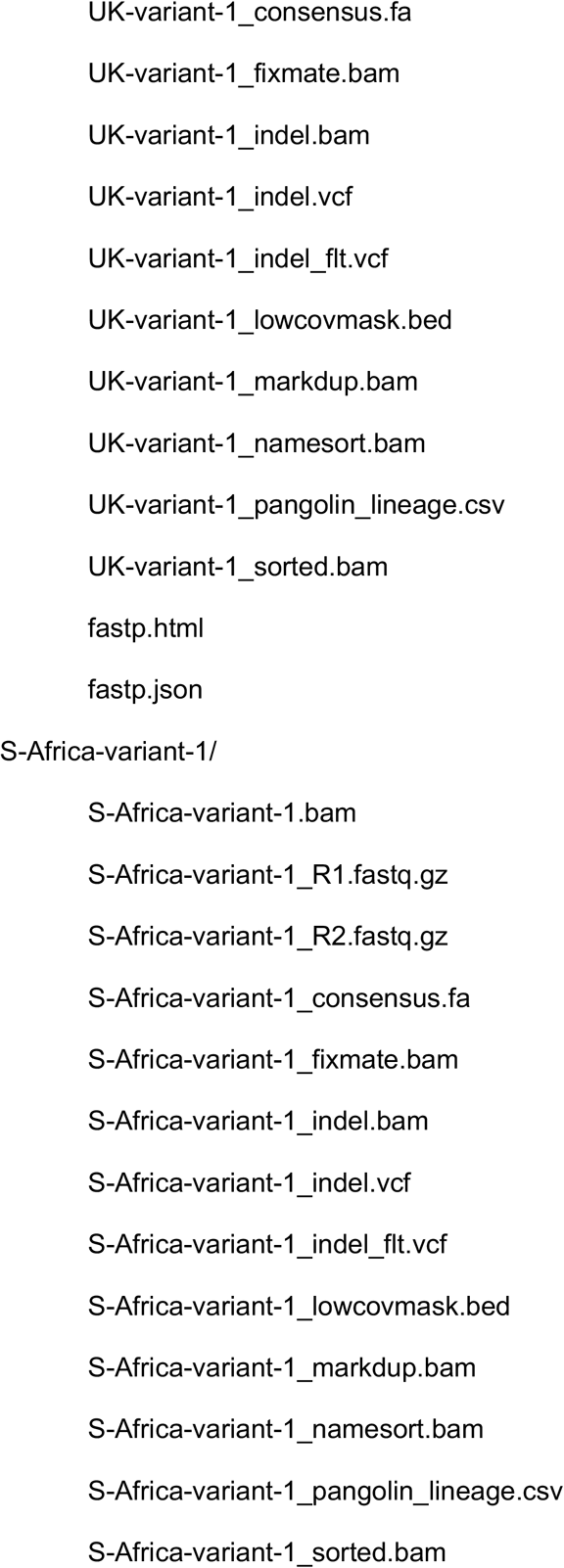

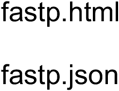

Each of the example UK variants should have been categorized as B.1.1.7 and the South African variants as B.1.351 (with pangoLEARN release 2021-02-06). It is important to note however, that as more sequences are uploaded and the pangolin lineage nomenclature updated, the assigned lineages may differ from the expected ones described in this paper. Regions with low coverages (with default setting under 30) are marked with the letter N during masking and represent gaps in the final consensus sequences.

*HaVoC* is comparable to alternative combinations of tools, e.g. Jovian and pangolin, in both speed and accuracy. These tools however operate separately, and as of publishing, there are no single public tools that can both perform a reference-based consensus assembly and a lineage identification in an easily accessible manner.

## Conclusions

Early detection and understanding of the potential impact of emerging variants of SARS-CoV-2 is of primary importance and can assist in more efficient surveillance and control of the disease. The likelihood of emergence of novel SARS-CoV-2 variants of concern is increased and accelerated by the high mutation rates typical in RNA viruses and the growing number of transmissions and infections both locally and globally.

With the rising number of variants detected worldwide and with many of them associated with increased transmissibility and lower vaccine efficacy, there is an emerging need for fast, efficient and reliable pipelines to help detect, identify and trace SARS-CoV-2 lineages. These pipelines should in addition be accessible to researchers who may not be familiar with utilizing complex bioinformatic tools or scripting pipelines.

Due to these challenges, we have developed *HaVoC*, a simple, reliable and user-friendly pipeline, which can be simply downloaded from our repository and run without being installed. All its dependencies can be installed via existing package managers, of which we recommend Bioconda. *HaVoC* could help in the current pandemic situation by detecting variants of concern in the sequencing centers and public health or other organisations currently running and tracing variants of concern worldwide. *HaVoC* is currently utilized for detecting and tracing SARS-CoV-2 variants of concern, mainly B.1.1.7, B1.351 and P.1, in Finland.

## Availability and requirements

*<u>Project name</u>: HaVoC* (Helsinki university Analyzer for Variants Of Concern)

*<u>Project home page:</u> https://www.helsinki.fi/en/projects/havoc and https://bitbucket.org/auto_cov_pipeline/havoc*

*<u>Operating system(s):</u> Linux, Mac*

*<u>Programming language:</u> Shell script*

*<u>Other requirements:</u> Trimmomatic or Fastp, BWA-MEM or Bowtie2, Samtools, BEDtools, BCFtools, Lowfreq and Pangolin.*

*<u>License:</u> GNU GPL*

*<u>Any restrictions to use by non-academics:</u> license needed*

## List of abbreviations

SARS-CoV-2: Severe acute respiratory syndrome coronavirus 2
COVID-19: Coronavirus disease 2019
*HaVoC*: Helsinki university Analyzer for Variants Of Concern

## Declarations

## Ethics approval and consent to participate

Not Applicable.

## Consent for publication

Not Applicable.

## Availability of data and materials

Publicly available at https://bitbucket.org/auto_cov_pipeline/havoc.

## Authors’ contributions

Conceptualization: PTN IP RK TS TSi OV. Development: PTN IP RK TS. Testing/Formal analysis: PTN IP RK TS. Funding acquisition: TSi OV. Investigation: PTN IP RK TS. Methodology: PTN IP RK TS. Project administration: RK TS OV. Resources: PTN RK IP TS TSi OV. Validation: PTN IP RK TS. Writing – original draft: PTN RK. Writing – review & editing: IP TS TSi OV.

## Acknowledgements

None.

## Authors’ information

None.

